# THE STRUCTURES OF MATURE AND IMMATURE ST. LOUIS ENCEPHALITIS VIRUS REVEAL CONSERVED HISTIDINE RESIDUES AFFECTING VIRUS FITNESS

**DOI:** 10.64898/2026.07.03.736192

**Authors:** Laís D. Coimbra, Samuel L. Guimarães, Luiza Leme, Alice Nagai, Marina A. Fontoura, Julia Rubiato, Leonardo C. Oliveira, Victória Bernardi, Guilherme Campos, Maurício L. Nogueira, Talita D. Melo-Hanchuk, Celso Benedetti, Rafael E. Marques

**Affiliations:** Brazilian Biosciences National Laboratory (LNBio), Brazilian Center for Research in Energy and Materials (CNPEM), Campinas, São Paulo, Brazil; Department of Genetics, Evolution, Microbiology and Immunology, Institute of Biology, University of Campinas (UNICAMP), Campinas, São Paulo, Brazil; Departamento de Doenças Dermatológicas, Infecciosas e Parasitárias, Faculdade de Medicina de São José do Rio Preto (FAMERP), São José do Rio Preto, São Paulo, Brazil; Departament of Pathology, University of Texas Medical Branch, Galveston, TX, USA

**Author notes:** Address correspondence to the last author,. These authors contributed equally to this article (Laís D. Coimbra, Samuel L. Guimarães).

## Abstract

Orthoflaviviruses undergo significant structural changes through maturation and infection, yet the molecular mechanisms remain incompletely understood. Using St. Louis encephalitis virus (SLEV) as a model, a reemerging mosquito-borne orthoflavivirus endemic in the Americas, we elucidated the structures of immature and mature SLEV particles at resolutions of 4.4 Å and 3.3 Å, respectively, using cryo-EM. SLEV is characterized by glycosylated E and prM proteins, the presence of lipid pockets, and is stabilized by an intricate network of inter- and intra-protein interactions between E and (pr)M across maturation stages. Several interactions were mediated by histidines that play different roles depending on SLEV maturation and pH. Non-lethal single mutations of H285R and H443R in E protein affect SLEV replication in mammalian and mosquito cell lines, and delay death in mouse models of infection. These histidine residues are conserved across orthoflaviviruses, illustrating the complexity of orthoflavivirus particles and indicating a possible strategy for attenuation.

## INTRODUCTION

St. Louis Encephalitis virus (SLEV, *Orthoflavivirus louisense*) is a reemerging arbovirus endemic in the Americas^1,2^. Similar to other arboviral pathogens such as Japanese Encephalitis virus (JEV) and West Nile virus (WNV), SLEV is maintained in a sylvatic cycle and is transmitted to humans by *Culex* mosquitoes^3–5^. SLEV infection in humans can cause manifestations ranging from an unspecific febrile disease to meningoencephalitis, which may result in sequelae or death^6,7^. The biology of SLEV is largely unknown, which has impaired the development of countermeasures against the disease. Nonetheless, mounting evidence of SLEV active circulation in the USA, Argentina, and Brazil^8–10^ indicates that SLEV poses a significant risk to public health.

SLEV is an enveloped single-stranded RNA orthoflavivirus. Orthoflaviviruses are known to generate mature and immature viral particles during replication: mature particles are infective and have ∼50 nm in diameter, while immature viral particles are slightly larger and not infective, but have been implicated in the development of severe forms of disease, notably in dengue^11^. Both forms are composed of structural proteins Capsid (C), (pre)Membrane [(pr)M] and Envelope (E), and maturation is achieved by host-mediated cleavage by furin and release of pr, leading to a rearrangement of M and E from a trimeric, spike-like conformation to M-E dimers organized sideways, decorating a smoother viral surface^12,13^. C is associated with the viral RNA in the interior of the viral particle. During infection, E, a class II fusion protein, engages with cellular receptors, resulting in orthoflavivirus internalization into endosomes^12^. Upon acidification, E proteins adopt an irreversible trimeric conformation that exposes the E protein fusion loop and promotes virus-host membrane fusion, resulting in the release of the viral genome in the target cell cytoplasm. The ∼11 kb SLEV genome encodes the three structural proteins and seven non-structural proteins (NS1, NS2a, NS2b, NS3, NS4a, NS4b, and NS5), all of which are translated by the host cell machinery and are involved in orthoflavivirus replication and assembly^7,14^. The molecular mechanisms underlying the significant structural changes observed in mature and immature orthoflavivirus particles are not completely understood.

Here we describe the structures of immature and mature SLEV particles obtained by cryo-EM at resolutions of 4.4 Å and 3.3 Å, respectively. SLEV presents a typical orthoflavivirus organization, with the presence of N-glycosylations in prM and E, and lipid pockets in the viral envelope. Comparison of immature and mature SLEV structures revealed critical interactions that stabilize protein conformation in different stages of virus maturation, several of which involve histidine residues (His/H). Non-lethal mutations on H285R and H443R in the E protein led to changes in SLEV replication in cell culture in a host-dependent manner, with single point-mutation H285R resulting in loss of fitness in mammalian (Vero CCL81) but not in mosquito cells (C6/36). SLEV infection of IFNAR^−/−^ and wild-type (WT) mice showed that both H285R and H443R resulted in attenuation in WT mice, with delayed mortality. E protein H285 and H443 are conserved across orthoflaviviruses, suggesting a rational approach for attenuation for other arboviral pathogens.

## METHODS

### Cell lines and Viruses

Vero CCL81, C6/36, and HeK293T cell lines were purchased from the BCRJ cell bank (Rio de Janeiro, RJ, Brazil). SLEV (GenBank accession number KM267635.1), isolate BeH355964, was provided by Dr. Luiz Tadeu Figueiredo (USP / Ribeirão Preto, Brazil). Vero and HEK293T cells were cultured in Dulbecco’s modified Eagle medium (DMEM; Cultilab, Brazil) supplemented with 10% of fetal bovine serum (FBS; Cultilab, Brazil) at 37 °C in 5% CO_2_ atmosphere. C6/36 cells were grown in Leibovitz medium (L-15; Cultilab, Brazil), supplemented with 1% of penicillin/streptomycin (Gibco, Thermo Fisher Scientific, USA) and 10% FBS, maintained at 28 °C without CO_2_.

### SLEV purification

SLEV BeH355964 purification was performed by infecting Vero CCL81 cells at a multiplicity of infection (MOI) of 10, followed by two supernatant collections at 48 h and 72 h post-infection. SLEV was precipitated from the total cell culture supernatant using 8% (w/v) PEG 8000 under gentle agitation overnight at 4 °C. Virus precipitates were collected by centrifugation at 16,800 × g for 50 min in an Avanti J-26S centrifuge (JLA-9.1000 Rotor), resuspended in NTE buffer (NaCl 120 mM, Tris 20 mM, EDTA 1 mM), centrifuged at 100,000 × g for 2 h (Optima XPN – 90, SW 41 Ti rotor) against a 22% sucrose cushion and again afterwards against a glycerol/ potassium tartrate gradient (10-35%) at 100,000 × g for 2 h. Purified SLEV bands were collected and concentrated using a 100 kDa cut-off filter (Amicon Ultra-2, Millipore). Purified SLEV samples were quantified using plaque-forming assays to determine the yield of infective viral particles and stocked at -80 °C until use.

### Negative staining and cryo-EM sample preparation

Purified SLEV samples were first evaluated by negative-stain transmission electron microscopy. Three microliters of sample were applied to glow-discharged Ultrathin Carbon Film on Lacey Carbon Support grids (#01824, TED PELLA) (1 min adsorption), stained twice with 2% (w/v) uranyl acetate (30 s incubation each), blotted with filter paper between each step, air-dried, and imaged on a JEM-1400 Plus transmission electron microscope (JEOL) operated at 80 kV and equipped with a 4k × 4k CMOS camera.

For cryo-EM, 3 µL of sample was applied to glow-discharged Quantifoil R2/1 grids (#662-200-CU, TED PELLA) and vitrified in liquid ethane using a Vitrobot Mark IV (Thermo Fisher Scientific). Blotting conditions were optimized empirically (force -4 to -5; blot time 3-5 s). Grids were screened on a Talos Arctica G2 microscope (Thermo Fisher Scientific) operated at 200 kV, and those displaying optimal ice thickness and particle distribution were stored in liquid nitrogen until data collection. All micrographs were collected at LNNano/CNPEM.

### Data collection and processing

Data were collected from two selected grids using *EPU* software on a Titan Krios G3i microscope (Thermo Fisher Scientific) operated at 300 kV and equipped with a Falcon 3EC direct electron detector. Movies were recorded in counting mode at 59,000× magnification (1.10745 Å/pixel), with a total dose of 30 e⁻/Å² distributed over 40 frames and a defocus range of -1.5 to -2.5 μm.

A total of 16,990 movies were imported into *SCIPION 3*, where all subsequent processing steps were performed. Motion correction, background subtraction, and CTF estimation were performed using *MotionCor2*, *Xmipp 3*, and *CTFFIND4*, respectively. After quality filtering, 16,563 micrographs were retained.

Manual particle picking was initially performed in *Xmipp 3* on a subset of micrographs to train the automated picking algorithm. Automated picking was subsequently applied to 10% of the dataset, followed by extraction and 2D classification in *RELION 4*. The best mature and immature 2D class averages were used as templates for reference-based autopicking across the full dataset. Particles were extracted and subjected to multiple rounds of 2D classification, after which mature and immature particles were separated based on morphology and processed independently.

Successive rounds of 2D classification, initial model generation, and 3D classification yielded homogeneous subsets of 50,475 immature and 49,349 mature particles. These were refined in *RELION 4* with imposed icosahedral symmetry, followed by CTF refinement, including corrections for beam tilt, higher-order aberrations, anisotropic magnification, astigmatism, and per-particle defocus. Final maps were obtained after additional refinement, masking, and post-processing. Global resolution was estimated using the gold-standard FSC 0.143 criterion, and local resolution was assessed in *RELION 4*. Parameters for cryo-EM data collection and processing, and model statistics are summarized in **Supplementary Table 1**.

### Localized reconstruction

Localized reconstructions of the asymmetric units (ASUs) were performed using *LocalRec* within the *SCIPION 3* framework. Sub-particle coordinates were defined in *UCSF Chimera 1.19* (CMM format) and used for extraction, followed by focused 3D classification, refinement, and post-processing in *RELION 4*. Global and local resolution were estimated from the final reconstructions.

A schematic overview of the single-particle analysis workflow, including representative 2D class averages, 3D classifications, and refined reconstructions, is provided in **Supplementary Fig. 3**.

### Model building and refinement

Atomic models were built from the ASU maps using *AlphaFold 3* models as initial templates. Models were fitted into the density maps in *UCSF Chimera 1.19* and refined through iterative cycles of manual adjustments in *Coot 0.9.8.95* and real-space refinement in *Phenix 2.0-5885*. Validation was performed with *MolProbity*, and structural figures were generated in *UCSF Chimera 1.19* or *ChimeraX 1.10.1*.

### Construction of recombinant viruses

Recombinant SLEV variants were generated using a modified circular polymerase extension reaction (CPER)-based strategy^15^. DNA constructs corresponding to the SLEV BeH355964 isolate were synthesized in the pcDNA 3.1 vector (Genscript) and used as templates to amplify the complete viral genome in three fragments: Fragment 1 = nt 1 to 2462, Fragment 2 = nt 2463 to 6702, and Fragment 3 = nt 6703 to 10939. Primers introduced a 40 bp overlap between adjacent fragments and the cloning vector. Ten full-length recombinant constructs were designed to encompass the complete SLEV genome sequence, including eight variations of fragment 1 with different mutations (**Supplementary tables 2 and 3**). The PJW377 vector, kindly provided by Dr. James Weger-Lucarelli, served as a CPER backbone and contains a CMV promoter immediately before the cloning site, and the HDV ribozyme sequence immediately after. Viral fragments were amplified, the vector linearized, and full-length recombinant genomes were assembled by CPER using Q5® High-Fidelity DNA Polymerase (New England Biolabs). CPER products were transfected into HEK293T cells using Lipofectamine 3000 (Invitrogen). Rescue efficiency was assessed by RT-qPCR using SYBR™ Green PCR Master Mix (Thermo Scientific) from viral RNA from culture supernatant 8 days post-transfection. The stability of recombinant viruses was assessed after two passages in Vero CCL81 cells by sequencing, using a MiSeq System (Illumina, USA). For both experiments, viral RNAs were extracted from culture supernatants with the QIAamp Viral RNA Mini Kit (Qiagen).

### *In vitro* SLEV replication curves

The replication fitness of SLEV BeH355964 and recombinant variants was evaluated using growth curves (SLEV WT; SLEV H81R; SLEV H398R; SLEV H81R/H398R; SLEV H285R; SLEV H285D; SLEV H443R; SLEV H443D). Vero CCL81 and C6/36 cells were plated in 96-well plates at densities of 5 × 10^3^ and 1 × 10^4^ cells/well, respectively. After 24 h, both cells were infected at an MOI of 0.1. Samples were collected at 12, 24, 48, and 72 h post-infection, and the viral titer was determined using a plaque-forming assay.

### Virus plaque-forming assay

Samples were serially diluted 1:10 in DMEM medium (0% FBS) and then added onto a monolayer of Vero CCL81 cells in 24-well plates and incubated for 1 h at 37 °C in 5% CO_2_. After this period, the inoculum was discarded, and the wells were covered with semi-solid medium containing 1% carboxymethylcellulose (CMC; Synth, São Paulo, Brazil) in 1:1 DMEM, 5% FBS, and 1% penicillin/streptomycin for 7 days. At the end, cells were fixed with 8% (w/v) paraformaldehyde (PFA; Sigma, St. Louis, MO, USA) and stained with 1% (w/v) methylene blue (Sigma, USA).

### Serial passage of SLEV in cell culture

SLEV evolution was evaluated using intra-host infection cycles in mosquito Aag2 and mammalian LLC-MK2 cell lines. Aag2 cell cultures were maintained at 28 °C in L15 media, and LLC-MK2 at 37 °C in MEM, both supplemented with 2% FBS. Viral RNA was extracted after each passage and sequenced using the MiSeq platform with the Illumina COVIDSeq library kit, applying a SLEV-specific primer set (**Supplementary table 4**). Data was analyzed using the *Geneious* software. Full SLEV genomes were obtained by both De Novo and Map to Reference approaches (to avoid bias during detection of nucleotide variations) and submitted to the variant call to identify mutations and frequency variation through passages.

### Mice

IFNAR^−/−^ mice (type I interferon receptor knockout) (B6.129S2-Ifnar1tm1Agt/Mmjax; JAX #010830) were provided by the animal facility LMEA at the Brazilian Biosciences National Laboratory (LNBio/CNPEM). C57BL/6J mice (JAX #000664) were purchased from the Multidisciplinary Center for Biological Research at UNICAMP. All *in vivo* procedures were conducted in a BSL-2 animal facility at LNBio/CNPEM. Animals were maintained under controlled temperature (21-24 °C) and a 12 h light/dark cycle, with *ad libitum* access to food and water.

### *In vivo* experiments

IFNAR^−/−^ and C57BL/6J mice, 8-10 weeks old, were infected with 10² PFU of SLEV WT, SLEV H285R, or SLEV H443R via subcutaneous (IFNAR^−/−^) or intracranial (C57BL/6J) route. Animals were monitored daily for 7 days for survival and weight analysis. Clinical signs, including ruffled fur, hunched posture, conjunctivitis, and asymmetrical paralysis of the lower limbs, were evaluated daily. Mice exhibiting signs of severe pain or distress were euthanized according to humane endpoint criteria and were considered dead for survival analysis. All procedures were performed according to Brazilian CONCEA regulations and approved by the local ethics committee (CEUA n°155).

### Statistical analysis

*In vitro* growth curves in Vero CCL81 and C6/36 cells were analyzed using one-way analysis of variance followed by the non-parametric Kruskal-Wallis test. A comparison of viral replication was performed between SLEV BeH355964 and recombinant variants of SLEV (SLEV WT; SLEV H81R; SLEV H398R; SLEV H81R/H398R; SLEV H285R; SLEV H285D; SLEV H443R; SLEV H443D) at each specific collection time (12, 24, 48, and 72 h).

*In vivo* survival curves were analyzed using the Log-Rank test for trend. Survival was compared between SLEV WT and SLEV H285R or SLEV H443R. *P*-values lower than 0.05 were considered significant. All statistical analyses were performed using *GraphPad Prism v10* (GraphPad Software, La Jolla, CA, USA).

## RESULTS AND DISCUSSION

### The Cryo-EM reconstructions of SLEV mature and immature viral particles

The concentration, viability, and purity of SLEV BeH355964 preparations were assessed by plaque-forming assays and negative-stain transmission electron microscopy. Micrographs revealed mostly spherical virions with an average diameter of ∼50 nm (**Supplementary Fig. 1A-C**), consistent with other orthoflaviviruses. Under negative-stain conditions, immature and mature particles were hardly discernible (**Supplementary Fig. 1B, C**). Conversely, Cryo-EM readily resolved immature, mature, and partially mature particles (**Supplementary Fig. 1D-E**). Particle abundance estimated from 2D class averages indicated an immature-to-mature ratio of approximately 1.35:1. Structural studies of orthoflaviviruses have consistently reported immature and mosaic particles in viral preparations^16–21^, reflecting incomplete or unsuccessful furin-mediated cleavage of prM before virus release. Our purified SLEV preparations contained a substantial proportion of immature particles when compared to other flavivirus^22^, possibly due to the high multiplicity of infection (MOI) used for SLEV propagation.

We leveraged this heterogeneity to resolve both immature and mature SLEV particles from the same cryo-EM dataset. Independent classification and refinement yielded full-particle reconstructions at global resolutions of 7.1 Å and 3.6 Å for immature and mature particles, respectively (**Fig. 1A, B, and Supplementary Fig. 2A, C**). Despite using more than 50,000 particles, the immature reconstruction remained poorly resolved because of pronounced positional variability (**Supplementary Video 1**)^23,24^. To overcome this limitation, localized reconstruction^25^ was used to isolate and independently align ASUs without imposing symmetry, improving the global resolution to 4.4 Å and 3.3 Å for the immature and mature particle, respectively (**Fig. 1C, D and Supplementary Fig. 2B, D**).

**Figure 1.**
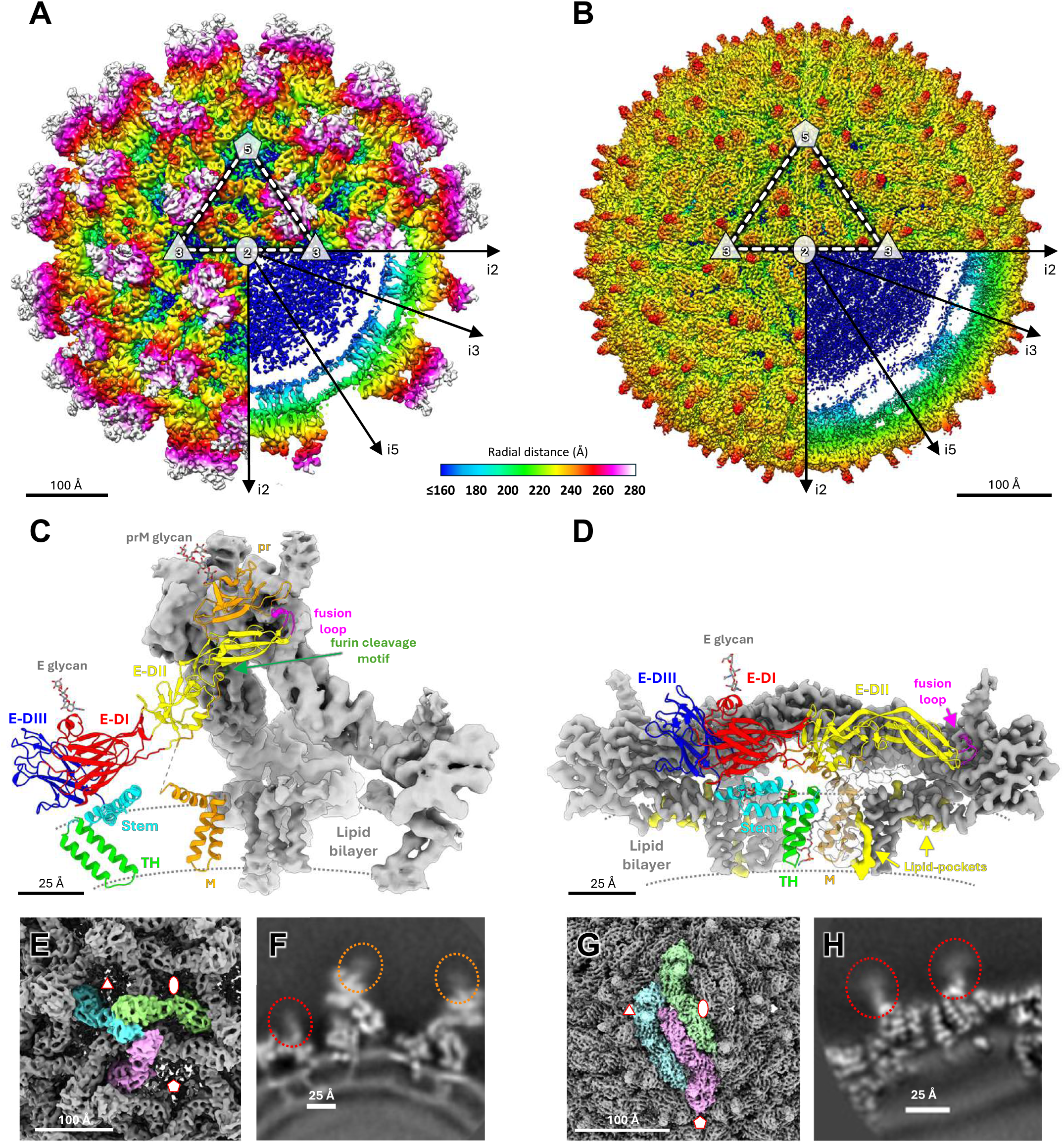
The Cryo-EM structures of immature and mature SLEV particles. Cryo-EM density maps of the full immature **(A)** and mature **(B)** particles were solved at 7.1 Å and 3.6 Å resolution, respectively, and colored by radial distance from the particle center. Icosahedral symmetry axes are indicated and dashed lines outline one theoretical ASU. The first three quadrants show the particle surface, whereas the fourth quadrant display a central slice to depict internal features. Locally reconstructed ASU from the immature **(C)** and mature **(D)** particles solved at 4.4 Å and 3.3 Å resolution, respectively, are also shown. One (pr)M-E heterodimer density is replaced by its atomic model shown as a cartoon. E domains and (pr)M are colored as follows: E-DI (red), E-DII (yellow), E-DIII (blue), Stem (cyan), transmembrane helices (TH, lime green), fusion loop (magenta), and (pr)M (orange); the furin cleavage motif is colored in green. The same color scheme is used throughout the other figures to facilitate structural interpretation. Glycans and conserved phospholipids are displayed as stick models. Grey dotted lines indicate the approximate lipid bilayer boundaries. In panel D, phospholipid densities are shown in yellow. The biological ASUs of immature **(E)** and mature **(G)** particles are shown as highlighted isosurfaces, with each (pr)M-E heterodimer colored cyan, green, or pink. The ⬠, △, and ⬯ symbols denote the i5-, i3-, and i2-fold symmetry axes, respectively. Panels **(F)** and **(H)** show slices through the locally reconstructed maps of the immature and mature particles. Dotted ellipses highlight glycans densities attached to E (red) and prM (orange). Scale bars are shown in Å.

### Structure of Immature SLEV

The immature particle is composed of 180 prM-E heterodimers anchored to the membrane by four transmembrane helices (TH) per heterodimer (two from each protein). These assemble into 60 trimeric spikes (prM_3_E_3_), each corresponding to a biological ASU (**Fig. 1E**), giving the particle its characteristic spiky appearance and a diameter of ∼57 nm. The core, which contains the viral RNA and C protein, does not exhibit icosahedral symmetry and cannot be resolved (**Fig. 1A**).

At the base of each pyramidal spike, the E domains-I and III (E-DI and E-DIII) lie adjacent to the membrane together with the Stem, E transmembrane helices (TH), and the M region of prM (**Fig. 1C**). E domain-II (E-DII) projects upward to form the central portion and base of the spike tip. At the tip, the β-sandwich pr domain caps the spike and shields the E-DII fusion loop from premature exposure. E-DI, E-DII, and E-DIII are predominantly β-stranded domains containing only a few short α-helices.

The pr and M domains are connected by an extended linker containing the furin cleavage motif that runs alongside E-DII toward the M helices (**Fig. 1C**). In our model, this linker is partially resolved, leaving a gap of 10 residues at the spike base near the membrane. This segment is also absent from most immature orthoflavivirus structures, suggesting increased conformational flexibility. At lower contour levels, a narrow density bridge spans this gap in one prM protomer, indicating the likely path of the unresolved segment (**Supplementary Fig. 4**). Although it cannot be modeled unambiguously, its position agrees with the arrangement reported for Binjari Virus (BinJV)^26^ and Tick-Borne Encephalitis Virus (TBEV)^27^. These observations support a model in which the immature orthoflavivirus spike is organized around a centrally positioned prM stem rather than a peripheral arrangement alongside the E TH^26^.

At the spike surface, both prM and E carry N-linked glycans, visible as diffuse but well-positioned cryo-EM densities (**Fig. 1F**). Each spike contains six glycans, one on each prM and E protomer (**Fig. 1C**), attached to N15 in the pr domain and N154 in E-DI. Up to five saccharide units could be modeled for prM glycans, whereas E glycans were less well resolved, likely due to steric constraints imposed by neighboring spikes. N-linked glycosylation at these positions is common among orthoflaviviruses, although occupancy and site conservation vary across species^28^. In prM, glycosylation has also been reported at N15 in WNV and JEV^29,30^, and the corresponding N-X-T/S motif is conserved among related encephalitogenic orthoflaviviruses (**Supplementary Fig. 5**). Alternative sites occur in other members, including N69 in Zika Virus (ZIKV) and Dengue virus (DENV)^31–34^. E glycosylation is more conserved, typically occurring at N154 or N153, when present, with few exceptions in DENV^35^ (**Supplementary Fig. 6**).

Mutagenesis studies in WNV, ZIKV, JEV, and TBEV demonstrated that these glycans are important for proper prM and E folding, viral assembly, and particle secretion^29,31,33,36^. A SLEV isolate (Tr 9464) lacking E glycosylation exhibited reduced viral yields and altered specific infectivity, consistent with morphogenetic defects caused by glycan loss^37^. N-linked glycans have also been implicated in cell attachment, infectivity, transmission, and virulence^29,30,34,38^. Altogether, the available evidence indicates that glycosylation of structural proteins is a multifaceted determinant of flavivirus assembly, host engagement, and pathogenesis. As proposed for other orthoflaviviruses, the immature SLEV assembly is a metastable architecture that precedes envelope rearrangements during maturation in the Trans-Golgi Network.

### Structure of Mature SLEV

Unlike the immature particle, the mature SLEV virion has a smooth, compact surface composed of 90 M_2_E_2_ dimers arranged in a herringbone pattern with icosahedral symmetry and a diameter of ∼50 nm (**Fig. 1B**). The ASU comprises three M and three E proteins, corresponding to one and a half M_2_E_2_ dimers (**Fig. 1G**). As in other orthoflaviviruses, this organization results from proteolytic removal of pr during maturation, allowing E-DII to move toward the lipid bilayer and bringing all three E domains into a more membrane-proximal arrangement (**Fig. 1D**). Simultaneously, the Stem and TH shift toward the ASU center, establishing stabilizing interactions with M. The remaining linker segment after the furin cleavage site and M packs beneath E-DII, further tightening the mature E shell. Glycans linked to E protein at N154 remain clearly visible, and three saccharide units could be modeled within the density (**Fig. 1D, H**). These glycans form the outermost features of the virion surface, positioned above the fusion loop of the adjacent E protomer within each dimer (**Fig. 1D**), consistent with other orthoflavivirus structures^18,21,27,34,35^.

### Conserved lipid-binding pockets in the mature SLEV virion

Extra well-defined densities were observed within the viral membrane boundaries, corresponding to three non-protein densities per M-E heterodimer (**Fig. 1D**). These densities could not be attributed to any structural protein and were therefore modeled as phosphatidylethanolamine molecules (3PE) occupying conserved lipid-binding pockets, as previously described in other orthoflaviviruses^24^.

Pockets 1 and 2 are located between the amphipathic helices of the Stem, whereas pocket 3 lies at the E-M transmembrane interface (**Fig. 2A**). Conserved residues stabilize the phospholipid head groups and portions of the acyl chains, indicating that these lipids are integral components of the mature envelope architecture (**Fig. 2B-D**).

**Figure 2.**
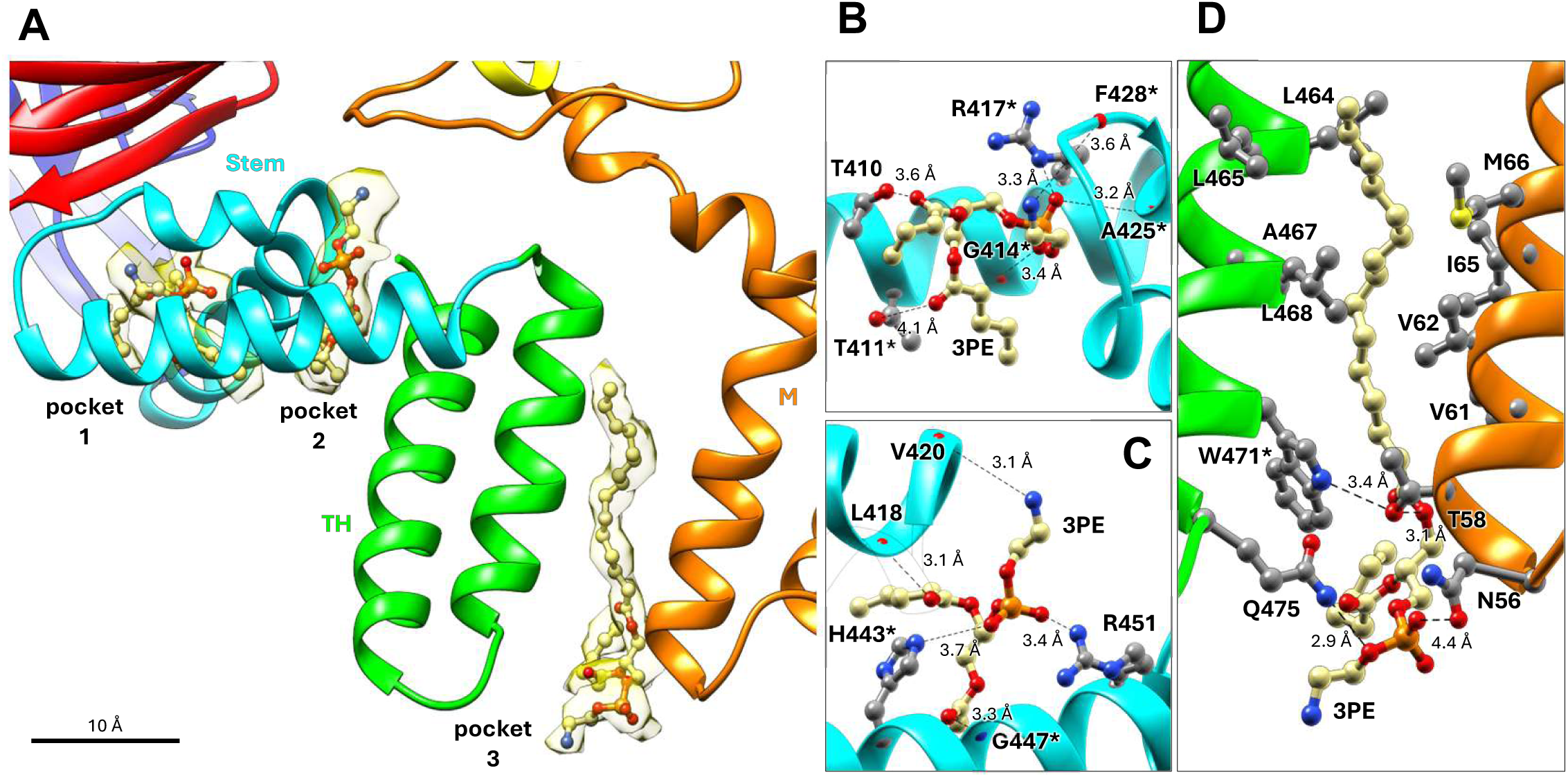
Conserved lipid-binding pockets in mature SLEV. **(A)** Overview of a single M-E heterodimer within the ASU, highlighting the three lipid-binding pockets. Pockets 1 and 2 are located between the amphipathic helices of the E Stem, whereas pocket 3 lies at the interface between the transmembrane helices of E and M. Modeled phosphatidylethanolamine molecules (3PE) are shown as ball-and-stick representations embedded within cryo-EM density (transparent yellow isosurface). Scale bar, 10 Å. **(B)** Pocket 1. The phospholipid head group is stabilized by R417* together with backbone atoms from G414*, A425*, and F428*. T410 and T411* are positioned near acyl-chain oxygens, providing additional polar contacts. The conserved glycine residue G414* contributes to formation of the lipid-binding cavity. **(C)** Pocket 2. The phosphate group is stabilized by R451 and the conserved H443*, whereas the phospholipid head group and acyl chains establish additional contacts with V420, L418, and G447*. The conserved G447* contributes to cavity formation for lipid accommodation. **(D)** Pocket 3 at the E-M transmembrane interface. The phospholipid head group is stabilized by Q475 from E and N56/T58 from M, while one acyl chain extends along the transmembrane helices and packs against surrounding hydrophobic residues. Distances and dashed lines indicate putative hydrogen bonds (Å). Conserved residues among orthoflaviviruses are marked with an asterisk (*).

Although the densities are well resolved, their quality does not allow unambiguous identification of the lipid species. Nevertheless, these pockets closely resemble those described in other orthoflaviviruses, where lipid cofactors stabilize the envelope architecture while maintaining the conformational flexibility required for maturation and infection. Consistent with their functional importance, disruption of analogous pockets in DENV and ZIKV markedly reduces infectivity^24^. The presence of conserved lipid-binding pockets in mature SLEV therefore suggests a similar structural role and highlights these sites as potential targets for broad-spectrum antiviral strategies.

### Structural roles of SLEV E histidines and pH-triggered conformational rearrangements

pH-induced rearrangements of the E protein occur near the p*Kₐ* of histidine, and protonation of one or more imidazole side chains has been proposed as a molecular switch driving structural transitions during virion maturation and membrane fusion^39,40^. The SLEV BeH355964 E protein contains 11 histidines, of which H144, H246, H320, and H443 are fully conserved across human-infecting orthoflaviviruses (**Supplementary Fig. 6**). We analyzed the structural environment of all histidines in immature and mature SLEV models and compared them with those in the fusogenic E conformation (PDB: 4FG0^41^) to assess their potential roles in particle stabilization and pH-dependent regulation of virus maturation and infectivity. Throughout this section, residues fully conserved among orthoflaviviruses pathogenic to humans are indicated by an asterisk (*).

### H144*, H152, and H320* stabilize the E-DI/E-DIII hinge in immature and mature E monomers

A network of polar interactions involving H144*, H152, and H320* stabilizes the E-DI/E-DIII intrachain interface in both immature and mature E monomers (**Fig. 3A, B**). Although the specific interactions differ between the two conformations, all three histidines remain associated with the hinge region, helping to maintain the relative positioning of E-DIII.

**Figure 3.**
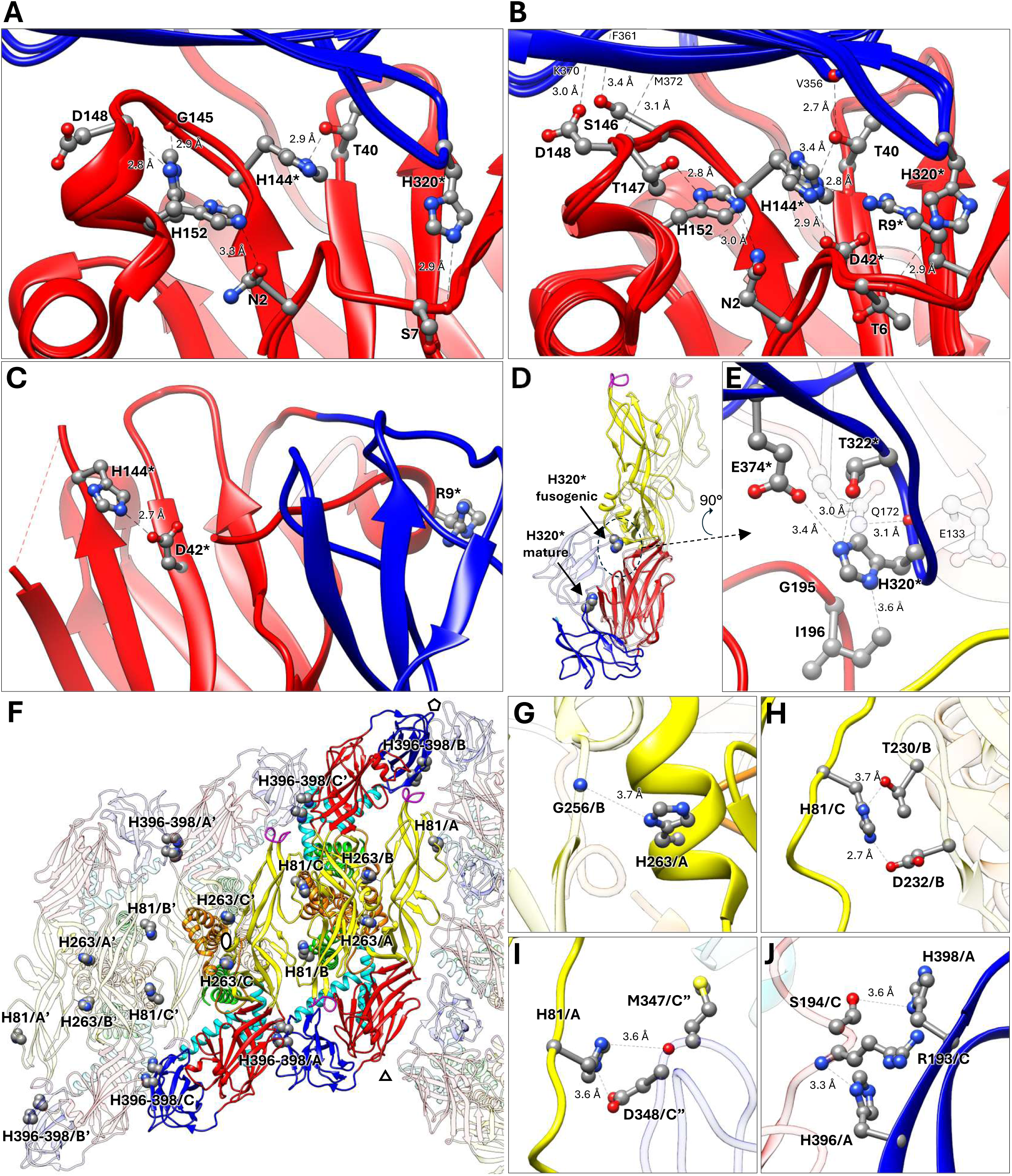
Histidine-mediated interactions in different conformations of the SLEV E protein and in envelope stabilization. Residues fully conserved among orthoflaviviruses infecting humans are indicated by an asterisk (*). **(A)** Close-up view of the E-DI/E-DIII interface in the immature SLEV E monomer, highlighting the interaction network formed by H144*, H152, and H320*. H144* interacts with T40. H152 contacts the backbone carbonyl groups of G145 and D148 (or N2 in an alternative protomer conformation). H320* interacts with the backbone carbonyl group of S7. **(B)** Corresponding view of the mature E conformation. H144* maintains contacts with T40 and additionally interacts with D42*. H152 interacts with T147 and/or N2. The H152-T147 interaction is uniquely conserved among JEV serogroup orthoflaviviruses. H320* remains associated with E-DI through contacts with the backbone carbonyl group of T6 and lies directly above R9*, forming, together with H144*, a compact basic cluster at the hinge region. Additional polar interactions involving T40-V356, S146-F361, T147-M372, and D148-K370 further stabilize the E-DI/E-DIII interface. **(C)** Fusogenic conformation showing retention of the H144*-D42* interaction after E-DIII rearrangement, the only his-mediated interaction preserved in all viral stages. The loop and short helix containing H152 and the glycosylated residue N154 are disordered and therefore absent from the atomic model. R9* is displaced away from the histidine cluster relative to the mature conformation. **(D)** Superposition of the mature E monomer (solid colors) and the fusogenic E conformation (faded colors; PDB ID: 4FG0), aligned using E-DI as a reference, illustrating the ∼32 Å displacement of H320* as E-DIII moves toward E-DII. **(E)** Interaction network adopted by H320* in the fusogenic conformation. The imidazole ring interacts with T322* and E374* within E-DIII and forms packing contacts with G195 and I196 from E-DI. The backbone carbonyl group of H320* is positioned near Q172 from a neighboring protomer, whereas the imidazole ring lies close to E133, a conserved residue in orthoflaviviruses, potentially forming an interprotomer electrostatic interaction. Secondary structure elements and residues from the second protomer in the fusogenic trimer are shown in semi-transparent representation. **(F)** Overview of the interaction network formed by H81, H263, and H396/H398 at intra- and interdimer interfaces in the mature virion. H263 stabilizes intradimer contacts, whereas H81 and H396/H398 connect neighboring dimers. The ⬠, △, and ⬯ symbols indicate the positions of the 5-, 3-, and 2-fold symmetry axes within the ASU. **(G)** Close-up of the intradimer interface showing interaction of H263 with G256 from the adjacent protomer. **(H-I)** Interdimer interactions mediated by H81. At internal interfaces within the same ASU **(H)**, H81 interacts with T230 and D232 from a neighboring protomer. At peripheral interfaces **(I)**, H81 instead contacts M347 and D348 of a protomer from a neighboring ASU, illustrating position-dependent interaction networks within the mature lattice. **(J)** Interdimer contacts involving H396 and H398. In chains A and C, H396 and H398 interact with R193 and S194 from adjacent dimers, whereas in chain B, both histidines are solvent-exposed. One or both His and R193 are present in most orthoflaviviruses. In panel **F**, faded colors denote protomers belonging to neighboring ASUs. In panels **G-J**, faded colors denote protomers other than the one containing the highlighted histidine.

Among these residues, H144* displays the most conserved interaction pattern, remaining associated with highly conserved residues within E-DI across all three conformations (**Fig. 3A-C**). H152 adopts a different interaction network after maturation but remains positioned within the hinge region (**Fig. 3A, B**). H152 and its interaction partner T147 are uniquely conserved among orthoflaviviruses from the JEV serogroup (**Supplementary Fig. 6**), suggesting that this interaction may represent a lineage-specific feature.

In both immature and mature conformations, H320* interacts with the backbone atoms from E-DI, while in the mature virion, it lies immediately adjacent to R9*, forming, together with H144*, a compact basic cluster at the E-DI/E-DIII hinge (**Fig. 3B**). Protonation of these two histidines under acidic conditions would increase the local positive charge density, potentially destabilizing this interface and facilitating the structural rearrangements required for membrane fusion.

The transition to the fusogenic conformation is accompanied by a major reorganization of this region. Whereas the H144*-D42* interaction is retained, the loop containing H152 and the glycosylated residue N154 becomes disordered and is not resolved (**Fig. 3C**). H320* undergoes the largest positional change, shifting approximately 32 Å as E-DIII moves toward E-DII (**Fig. 3D**). Consequently, the mature basic cluster is disrupted and H320* becomes incorporated into a new conserved interaction network with T322* and E374* that likely contributes to stabilization of the fusogenic trimer (**Fig. 3E**). This interpretation is supported by previous studies showing that the equivalent histidine in TBEV functions as a pH sensor for membrane fusion, whereas H320* and E374* are required for virus entry and particle release in JEV, respectively^40^.

Together, the maturation-dependent rearrangements of R9*, H144*, and H320* support a model in which a conserved electrostatic cluster at the E-DI/E-DIII hinge promotes pH-triggered destabilization of the mature envelope and transition to the fusogenic conformation.

### H81, H263, H396, and H398 mediate intra- and interdimer interactions that connect E dimers in the mature SLEV

H81, H263, H396, and H398 are generally less conserved across orthoflaviviruses (**Supplementary Fig. 6**) and have not been previously implicated in major conformational rearrangements. Nevertheless, these residues contribute to stabilization of the mature envelope by connecting neighboring E protomers and dimers (**Fig. 3F**). H263 participates in intradimer contacts, stabilizing the interface between protomers within the same E dimer (**Fig. 3G**). Both H263 and its interaction partner G256 are highly conserved across orthoflaviviruses, indicating strong structural constraint at this interface.

In contrast, H81, H396, and H398 contribute primarily to interdimer contacts that connect adjacent E dimers within the mature lattice (**Fig. 3F, H-J**). H81 adopts distinct interaction patterns depending on its position within the ASU (**Fig. 3H, I**), whereas H396 and H398 participate in a conserved interaction network involving residues from neighboring dimers (**Fig. 3J**). Although the specific residues vary among orthoflaviviruses, the overall electrostatic character of these interfaces is generally preserved (**Supplementary Fig. 6**), suggesting conservation of function rather than strict conservation of sequence.

Among these residues, H396 and H398 are the strongest candidates for pH-dependent destabilization of interdimer contacts, as protonation would increase the positive charge density near R193 (**Fig. 3J**), potentially generating electrostatic repulsion that weakens interdimer contacts and facilitates envelope reorganization leading to membrane fusion.

Consistent with a structural role in the mature lattice, the interdimer contacts formed by H81, H396, and H398 are absent in the immature and fusogenic conformations, where these residues are largely solvent-exposed. In the fusogenic trimer, however, H81 becomes buried and positioned near D95 from a neighboring protomer, where it likely contributes to stabilization of the fusogenic architecture^41,42^.

Together, these residues form an extended interprotomer network that acts as a molecular zipper, locking adjacent E dimers within the mature envelope lattice.

### H285* and H443* coordinate rearrangements at the E-DI/Stem interface during maturation

In immature prM-E trimers, H285* and H443* participate in distinct interaction networks (**Fig. 4A**). Following the large-scale rearrangements associated with virion maturation, these interactions are extensively reorganized, bringing both histidines into a common structural region, maintaining the link between E-DI and the Stem (**Fig. 4B**).

**Figure 4.**
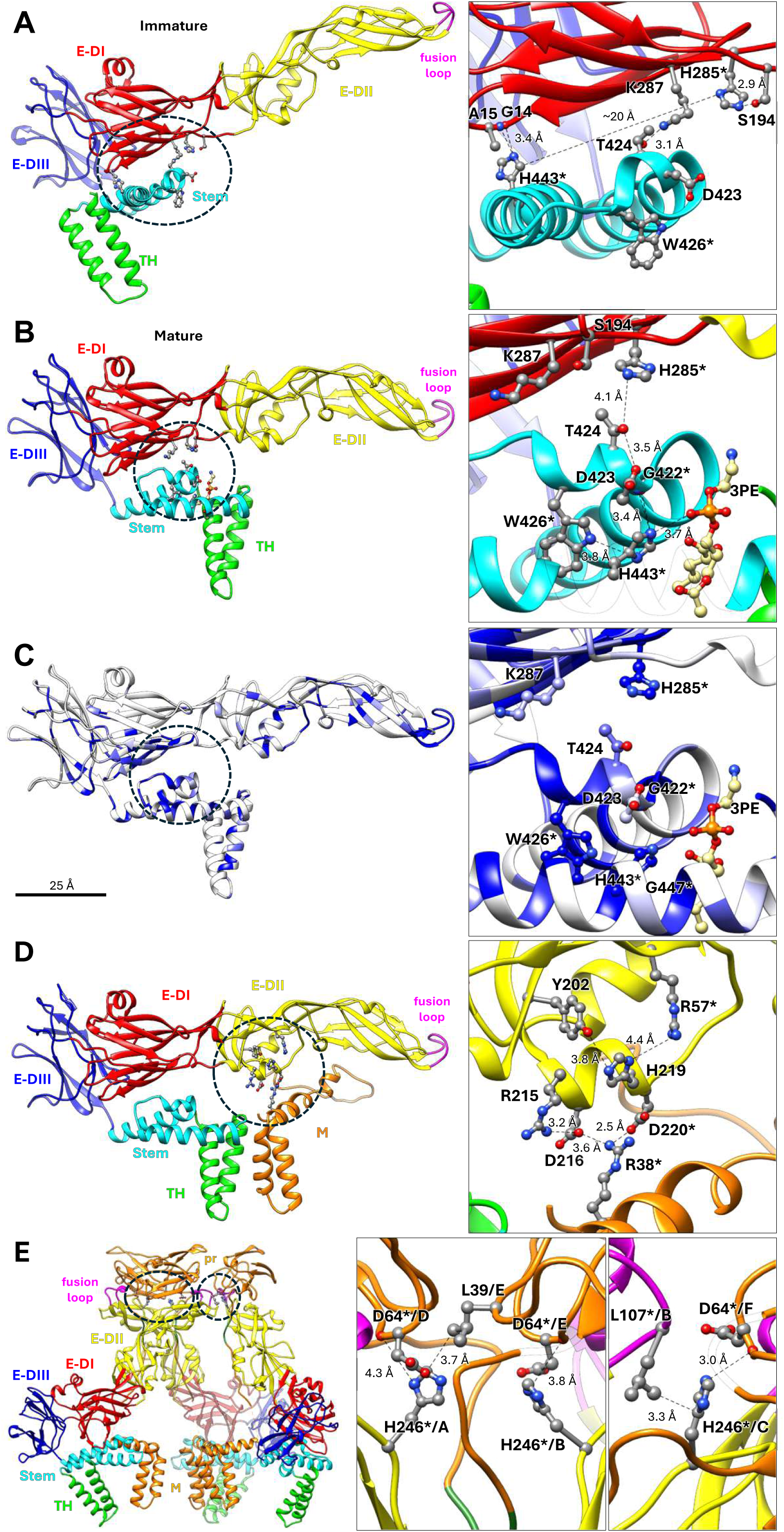
Histidine interaction networks involved in E-DI/Stem stabilization and E-(pr)M interactions. Residues fully conserved among human-infecting orthoflaviviruses are indicated by an asterisk (*). **(A)** Structural view of the immature E protomer. H285* forms a hydrogen bond with S194 in E-DI, whereas H443* from the Stem interacts with the backbone atoms of G14/A15 in E-DI. In this conformation, the two histidines are separated by ∼20 Å and participate in independent interaction networks. **(B)** Mature E protomer showing reorganization of the E-DI/Stem interface. H285* becomes positioned near T424, while H443* forms an extended network of polar interactions involving G422*, D423, W426*, and the head group of a phosphatidylethanolamine molecule (3PE) occupying lipid-binding pocket 2. D423 also interacts with T424, linking the Stem helices to E-DI through a conserved polar network. **(C)** Conservation mapped onto the E protomer. Conservation across human-infecting orthoflaviviruses is shown from white (≤70%) to blue (100%). Insets highlight residues involved in the H285*-T424-D423-G422*-H443*-W426* network and the architecture of lipid-binding pocket 2. The E-DI/Stem interface and Stem amphipathic helices correspond to some of the most conserved regions of the E protein, together with the fusion loop at the tip of E-DII. **(D)** Structural environment of H219 in the mature E protein. H219 is positioned within a conserved electrostatic network located at the E-M interface. Surrounding very conserved residues include R57*, Y202, R215, D216, D220*, and R38* from M. H219 does not directly participate in the principal salt-bridge network, it forms a hydrogen bond with Y202 and is located adjacent to residues that stabilize contacts between E-DII and the M amphipathic helix. Protonation of H219 may create a competition with R38* from M for salt-bridge with D220*, weakening interactions between E and M. **(E)** Structural environment of H246* in the immature prM-E spike. H246* lies near the fusion loop and participates in distinct interactions in each protomer of the trimeric spike. In chain A, H246* is positioned near D64* from its associated prM molecule and L39 from a neighboring prM-E heterodimer. In chain B, H246* interacts with D64* from its associated prM molecule. In chain C, H246* contacts both D64* from prM and L107* from a neighboring E protomer. D64* and L107* are fully conserved among orthoflaviviruses, whereas position 39 is occupied predominantly by hydrophobic residues. Owing to the resolution of the immature reconstruction, these interactions should be interpreted as structurally plausible. Insets show magnified views of the circled regions. Distances are indicated in Å.

In the mature conformation, H285* becomes associated with residues from the Stem, whereas H443* participates in an extended polar network that also contributes to the architecture of lipid-binding pocket 2 (**Fig. 4B**). Together with G422*, D423, T424, and W426*, these residues stabilize the membrane-proximal positioning of the Stem helices relative to E-DI.

The residues forming this network are among the most conserved region in orthoflavivirus E proteins (**Supplementary Fig. 6; Fig. 4C**). H285*, G422*, W426*, and H443* are fully conserved, whereas D423 and T424 retain chemically similar substitutions, indicating strong evolutionary constraints on this interface.

Consistent with this structural role, H443* appears to stabilize both immature and mature conformations through distinct interaction patterns, and mutations affecting this region severely impair viral viability^24^. H285* likewise contributes to stabilization of the mature E architecture by helping anchor E-DI against the Stem.

Together, these observations suggest that H285* and H443* participate in a conserved interaction network that couples stabilization of the mature E-DI/Stem interface with the structural organization of lipid-binding pocket 2. Because this network is absent in immature trimers and reorganized during maturation, these residues are also plausible contributors to pH-dependent envelope rearrangements.

### H219 and H246* contribute with distinct structural roles rather than acting as canonical pH sensors

H219, located in E-DII, is conserved in several orthoflaviviruses (**Supplementary Fig. 6**). In the mature virion, it occupies a conserved electrostatic environment close to the E-M interface (**Fig. 4D**). Although not directly involved in the main interaction network, protonation of H219 could alter the local electrostatic balance and weaken E-M contacts. Consistent with this interpretation, the surrounding network is extensively reorganized during maturation and absent in the fusogenic conformation. Thus, H219 may act as a local electrostatic modulator that indirectly facilitates E-M dissociation for fusion.

H246*, also located in E-DII, is solvent-exposed in mature and fusogenic conformations, while in the immature particle it participates in interactions involving neighboring E protomers and the pr domain (**Fig. 4E**). The conservation of residues surrounding H246* and the drastic decreased secretion caused by mutation of the equivalent residue in DENV2 (H244A), partially rescued by neutralization of the secretory pathway pH, suggest that H246* contributes to stabilization and proper maturation of the immature prM-E spike^43,44^.

### H285 and H443 are essential for SLEV viability and replication fitness in cell culture

According to our structural analysis, H81 and H398 are important for ‘glueing’ E dimers together in the mature conformation of SLEV, while the H285-H443 network of interactions plays a key role in the transition between the trimer-dimer conformations of E found in immature and mature SLEV particles, respectively. To identify residues critical for SLEV virion assembly and replication, we generated recombinant variants targeting these four His residues (H81, H285, H398, and H443) predicted to play roles in viral structural stability (**Fig. 5A**). In addition to a recombinant WT clone, seven constructs (H81R, H81R/H398R, H285D, H285R, H398R, H443D, H443R) harboring His-to-Arg or His-to-Asp substitutions were generated using the CPER method. Transfection of HEK293T cells with all constructs yielded detectable viral RNA by RT-qPCR at day 8 post-transfection (**Supplementary Fig. 7**), confirming successful genome transfection in all variants. However, infectious virus rescue was selective: five mutants (H81R, H398R, H81R/H398R, H285R and H443R) produced infectious particles, while H285D and H443D yielded no infectious virus despite comparable RNA levels, indicating that Asp substitutions ablate virus viability (**Fig. 5B**). To assess virus replication fitness in cell culture, replication curves were performed in mammalian (Vero CCL81) and insect (C6/36) cells at a MOI 0.1 (**Fig. 5C, D**). The WT recombinant SLEV replication curve was similar to the parental SLEV BeH355964 in both cell lines, though with slightly lower peak viral titer for the parental strain in C6/36 cells. H81R, H398R, and H81R/H398R recombinant viruses exhibited WT-like replication dynamics in Vero CCL81 cells (10⁶-10⁷ PFU/mL by 72 hpi). In contrast, H285R and H443R mutant viruses showed 1-2 log_10_ reduction in viral loads in the mammalian cell line, with delayed growth rates and lower plateau titers (**Fig. 5C**). Notably, all mutant viruses presented reduced replication dynamics in C6/36 cells, except for SLEV H285R, which performed similarly to WT SLEV (**Fig. 5D**).

**Figure 5.**
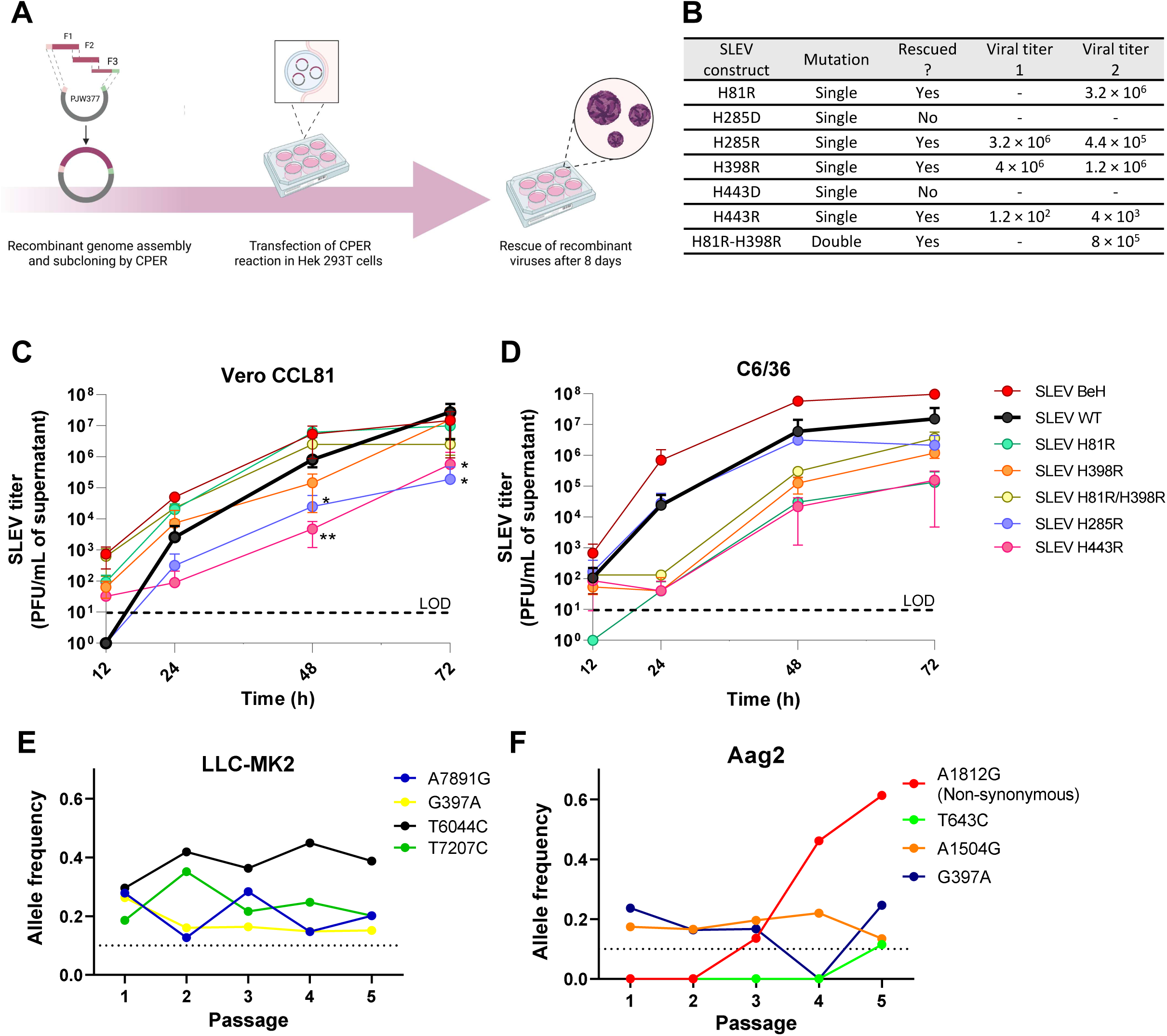
Mutations of E protein histidines affect SLEV fitness in cell culture. **(A)** Schematic workflow applied in the production of recombinant SLEV mutants by circular polymerase extension reaction (CPER) followed by virus rescue in 293T cells. **(B)** Summary of the E-protein histidine mutants generated and rescued after passage in VERO CCL81 cells in two independent experiments. Viral titers quantified by plaque lysis assay and expressed in PFU/mL. **(C)** Growth curves of SLEV mutants at an MOI of 0.1 in VERO CCL81 mammalian cells quantified by plaque-forming assay. **(D)** Growth curves of SLEV mutants at an MOI of 0.1 in C6/36 insect cells quantified by plaque-forming assay. **(E)** SLEV allele frequencies during 5 serial passages in LLC-MK2 cells. **(F)** SLEV allele frequencies along 5 passages in Aag2 cells. Data show the genetic stability and replicative fitness of the SLEV mutants across mammalian and mosquito cell environments.

Separately, we assessed SLEV evolution in cell culture, through serial passage in mammalian LLC-MK2 or mosquito Aag2 cells. Serial passage in LLC-MK2 cells resulted in the accumulation of 4 synonymous genetic mutations (A7891G, G397A, T6044C, T7207C) with stable frequencies throughout the experiment (**Fig. 5E**). In contrast, in Aag2 cells, we observed the emergence of a nonsynonymous mutation in A1812G in the E protein, which translates to H285R, together with three synonymous mutations (T643C, A1504G, G397A) (**Fig. 5F**). Successive SLEV passages led to a higher frequency of mutation H285R, rising from undetectable to 61% in the population by the third passage.

These results indicate that several His residues from E are important for SLEV biology, affecting viral replication in cell culture, but this effect varies between mammalian and insect hosts. While H443R was detrimental to SLEV in both hosts, H285R was detrimental in mammalian cells but not in insect cells and was spontaneously selected by serial passage in that cell type. Recombinant viruses were sequence-confirmed after two passages, but additional passages are required to determine the genetic stability of H285R and H443R. Altogether, our findings support key roles performed by His residues in SLEV E protein and indicate that their effects on viral fitness in cell culture are conditioned by the host, likely reflecting differences in secretory pathways, lipid composition and temperature between mammalian and mosquito cells.

### H285 and H443 affect SLEV pathogenicity in mouse models

Because recombinant SLEV bearing H285D and H443D could not be rescued from cell culture, and H285R and H443R were attenuated in mammalian cells, we hypothesized that mutated H285 and H443 might reduce SLEV pathogenicity *in vivo*. We used two highly susceptible mouse models of SLEV infection: intracranially-inoculated immunocompetent adult C57BL/6 mice and the Type I Interferon receptor-deficient IFNAR^−/−^ mice (**Fig. 6A-D**)^7,45^. Both models are lethal and characterized by an acute onset of severe disease, resulting in weight loss and death within 7 days post-infection (p.i.) with WT SLEV (**Fig. 6**). Infection with recombinant SLEV bearing the single-point mutation H285R resulted in delayed weight loss and death in IFNAR^−/−^ mice (**Fig. 6B, C**) and in a delayed death in C57BL/6 mice (**Fig. 6E, F**), while infection with SLEV H443R induced weight loss and death similar to WT SLEV in the IFNAR^−/−^ mice, but also delayed the onset of death in C57BL/6 mice (**Fig. 6E, F**). Altogether, these results indicate that mutations H285R and H443R moderately reduce SLEV pathogenicity *in vivo*, corroborating our previous findings in mammalian cell cultures. Considering H285 and H443 are conserved in orthoflaviviruses, such mutations could be explored in the rational design of attenuated strains for various applications for SLEV and related arboviral pathogens.

**Figure 6.**
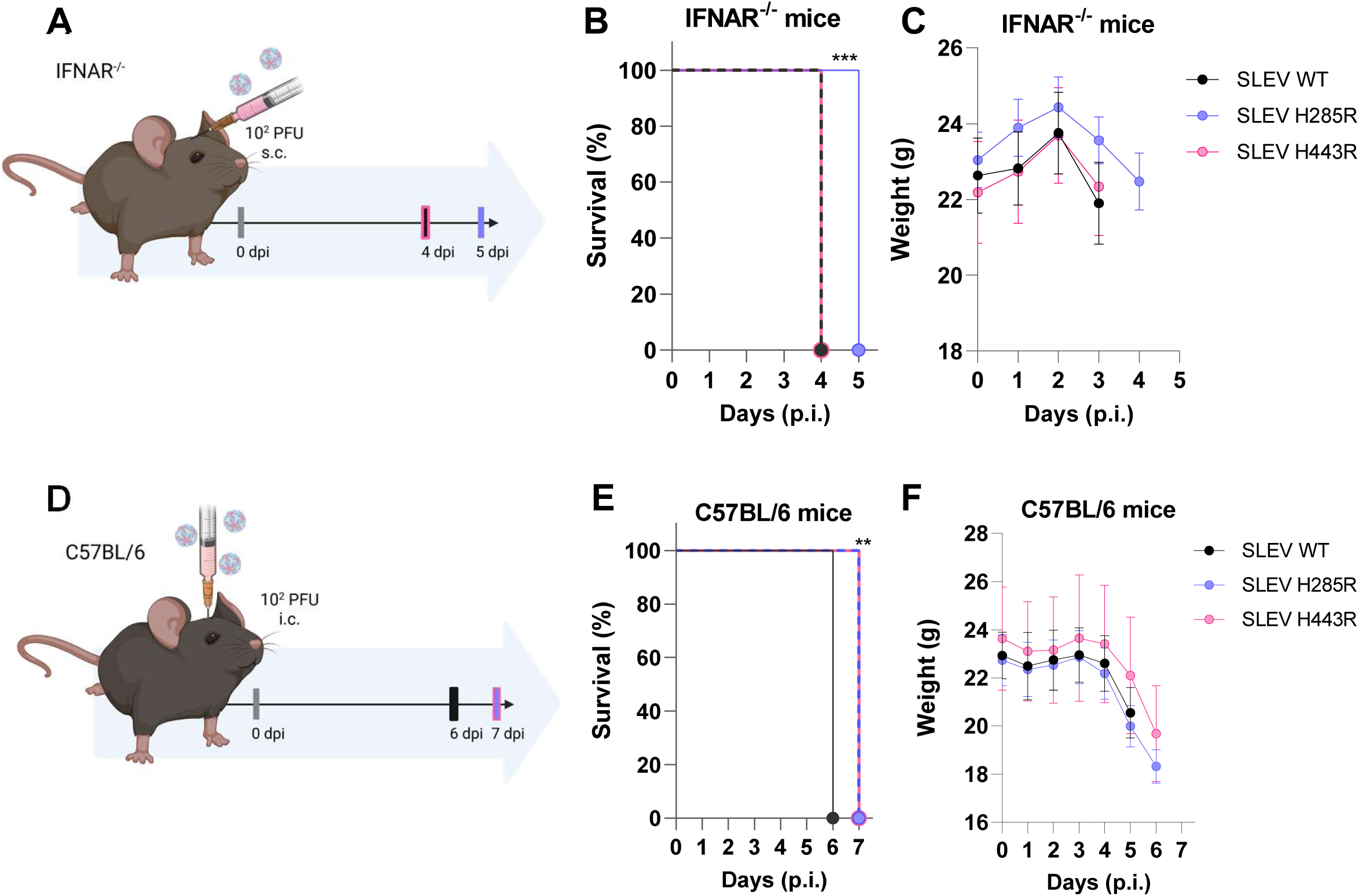
His285 and His443 affect SLEV pathogenicity in mouse models. Adult mice were infected with 10^2^ PFU of SLEV WT (black), SLEV H285R (blue) or SLEV H443R (pink) and monitored daily. Mortality occurred when mice either succumbed to disease or were humanely euthanized. **(A)** Scheme representing the experimental design of subcutaneous SLEV infection in IFNAR^−/−^mice. **(B)** Kaplan-Meier survival curves of SLEV-infected IFNAR^−/−^ mice. **(C)** Weight tracking throughout the five days of survival experiment. **(D)** Scheme representing the experimental design of intracranial SLEV infection in C57BL/6J mice. **(E)** Survival curves of C57BL/6 mice. **(F)** Weight loss throughout the seven days of experiment. Log-rank test for trend (**p<0.01; ***p<0.001).

## Supporting information

Supplemental data sets

## DATA AVAILABILITY

The cryo-EM reconstructions obtained in this work (Full immature, Full mature, ASU immature, and ASU mature) are available in the EMDB under accession codes EMD-77127, EMD-77122, EMD-77179, and EMD-77099, respectively. The models corresponding to the ASU of immature and mature particles are available in the PDB under accession codes 35TE and 13JV, respectively. Raw experimental data can be provided upon request.

## ACKNOWLEDMENTS

We would like to thank Mr. Valber Ferreira for his technical support. We would like to thank Dr. Rodrigo Portugal and the entire team at the Electron Microscopy Facility at LNNano (proposals 20210252, 20221006, 20230467, and 20233626). We thank the LSB and LMEA (proposal 20262655) facilities at LNBio for access to the necessary infrastructure and resources. We would like to thank Prof. James Weger-Lucarelli for supporting the construction of the recombinant viruses. Finally, we thank Douglas Paixão and Dr. Douglas Adamoski for their support on genome sequencing.

## FINANCIAL SUPPORT

This work was supported by the Brazilian Ministry of Science and Technology and by FAPESP (2021/05519-0, 2023/10771-6,). This work was also supported by INCT-One Health and INCT-Dengue. MLN and REM are CNPq research fellows.

## AUTHOR CONTRIBUTIONS

LDC, SLG, LL, AN, MAF, JR, LCO, VB, GC, and TDMH performed experiments. LDC, SLG, and VB were involved in data processing. LDC, SLG, MLN, GC, TDMH, CB, and REM analyzed and interpreted data. CB and REM conceived the study. All authors were involved in manuscript writing and revision.

## COMPETING INTERESTS

Authors declare no competing interests.

## Notes

### Competing Interest Statement

The authors have declared no competing interest.

