## Supplemental data sets for "THE STRUCTURES OF MATURE AND IMMATURE ST. LOUIS ENCEPHALITIS VIRUS REVEAL CONSERVED HISTIDINE RESIDUES AFFECTING VIRUS FITNESS"

A

| Sample | Total viral load (PFU) |
|---|---|
| Infective SLEV particles recovered in supernatant of Vero infected cells | $3 \times 10^{10}$ |
| Infective SLEV particles preciptated by PEG 8000 | $1 \times 10^{10}$ |
| Infective SLEV particles purified in Sucrose cushion | $4 \times 10^{9}$ |
| Infective SLEV particles purified in tartrate gradiente | $7 \times 10^{8}$ |
| Infective SLEV particles after concentration in 100 kDa cut-off filter | $3 \times 10^{9}$ |

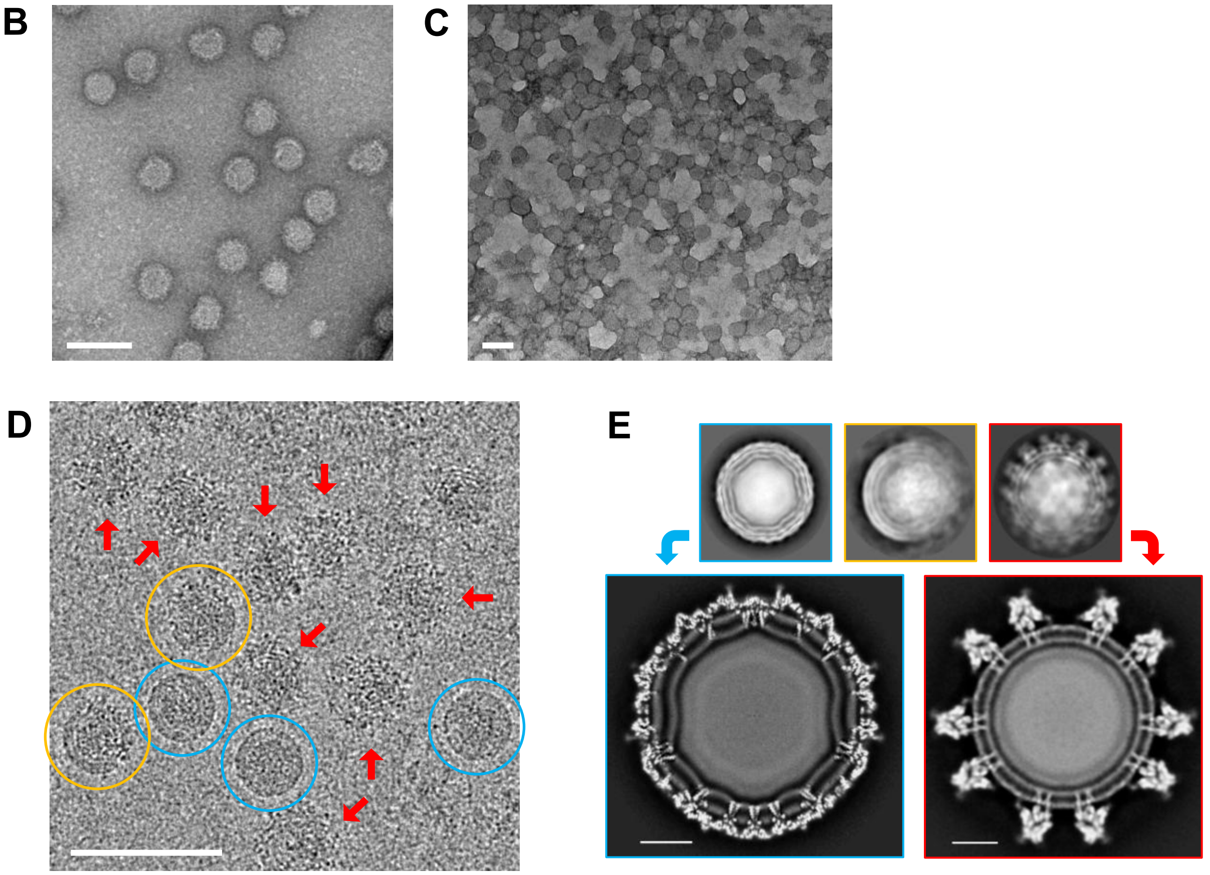


**Supplementary Figure 1 – Purification, negative-stain screening, and Cryo-EM identification of mature, immature, and partially mature SLEV particles. (A)** Viral recovery throughout the purification procedures, expressed as plaque-forming units (PFU) measured in the supernatant after each purification step. **(B)** Negative-stain transmission electron micrograph of the purified virus collected from the final purification band, showing predominantly spherical particles with diameters of approximately 50 nm. Under negative-stain conditions, mature and immature particles cannot be readily distinguished based solely on morphology. **(C)** Negative-stain micrograph of the concentrated purified sample, revealing a high density of viral particles suitable for subsequent cryo-EM grid preparation. **(D)** Representative cryo-EM micrograph of the concentrated sample embedded in vitreous ice. Mature particles (blue circles) display smooth outer contours, whereas immature particles (red arrows) exhibit diffuse, spiky edges resulting from the protruding prM-E spikes. Partially mature particles (yellow circles) display both smooth and spiky regions, consistent with incomplete maturation. **(E)** Top: representative 2D class averages corresponding to mature (blue), partially mature (yellow), and immature (red) particles. Bottom: central slices through the refined cryo-EM reconstructions of the mature (left) and immature (right) particles, highlighting the distinct envelope architectures of the two maturation states. Scale bars: 100 nm in B-D, 10 nm in E.

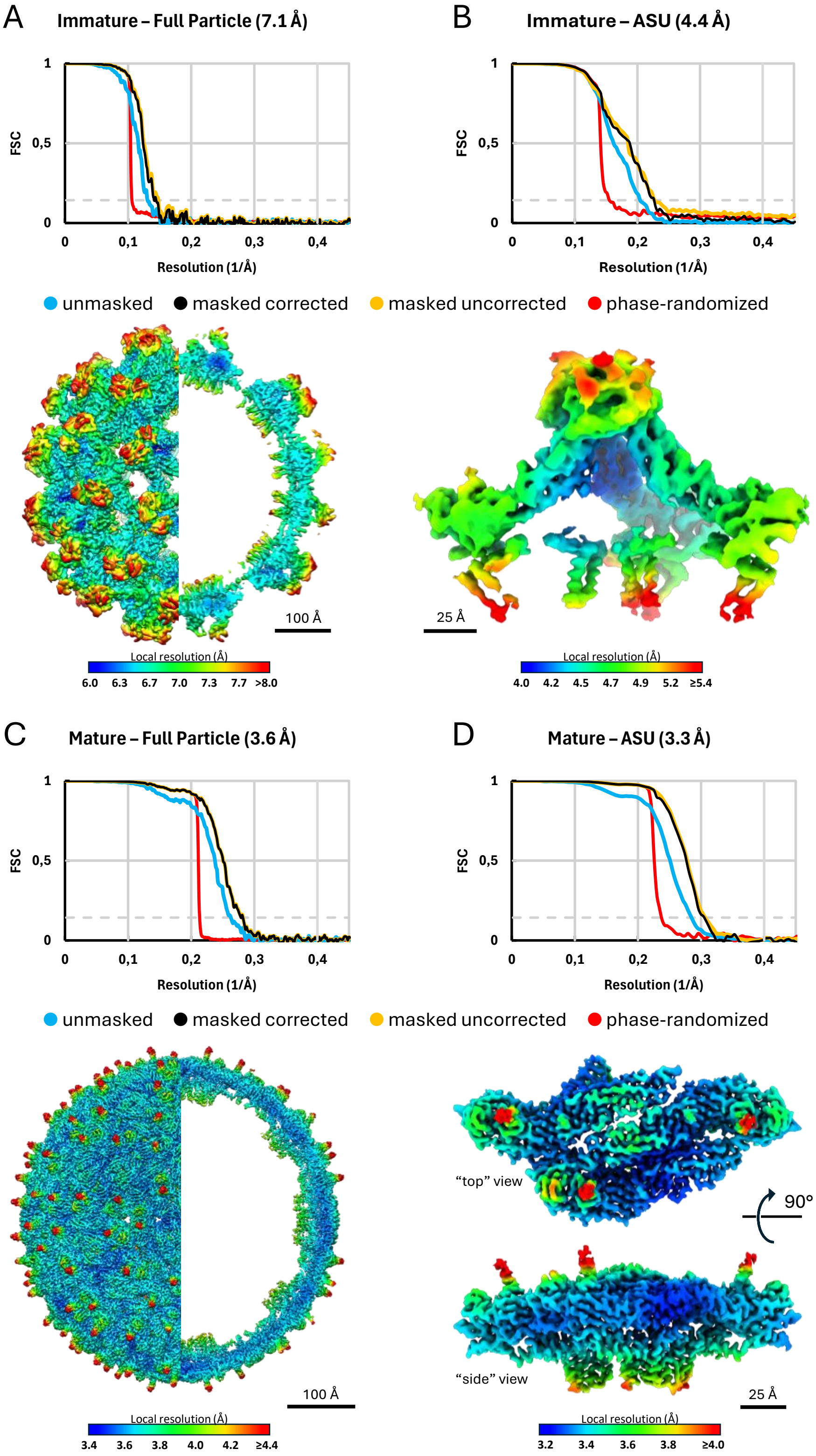


**Supplementary Figure 2 – Fourier shell correlation (FSC) and local resolution analysis of immature and mature SLEV reconstructions.** FSC curves of the full immature SLEV particle reconstruction **(A)** and the full mature SLEV particle reconstruction **(B)** are shown. **(C)** and **(D)** show FSC curves of the locally reconstructed ASU from the immature and mature particles, respectively. In all panels, masked (corrected and uncorrected), unmasked, and phase-randomized FSC curves are shown as indicated. The dashed line represents the 0.143 criterion used to estimate the global resolution. Below each FSC plot, the corresponding cryo-EM density map is shown as an isosurface representation colored according to the local resolution, as indicated by the color scales. For the full-particle reconstructions, the left half displays the external isosurface representation of the particle, while the right half presents a central cross-section to facilitate visualization of internal structural elements. Scale bars are shown in each panel (Å).

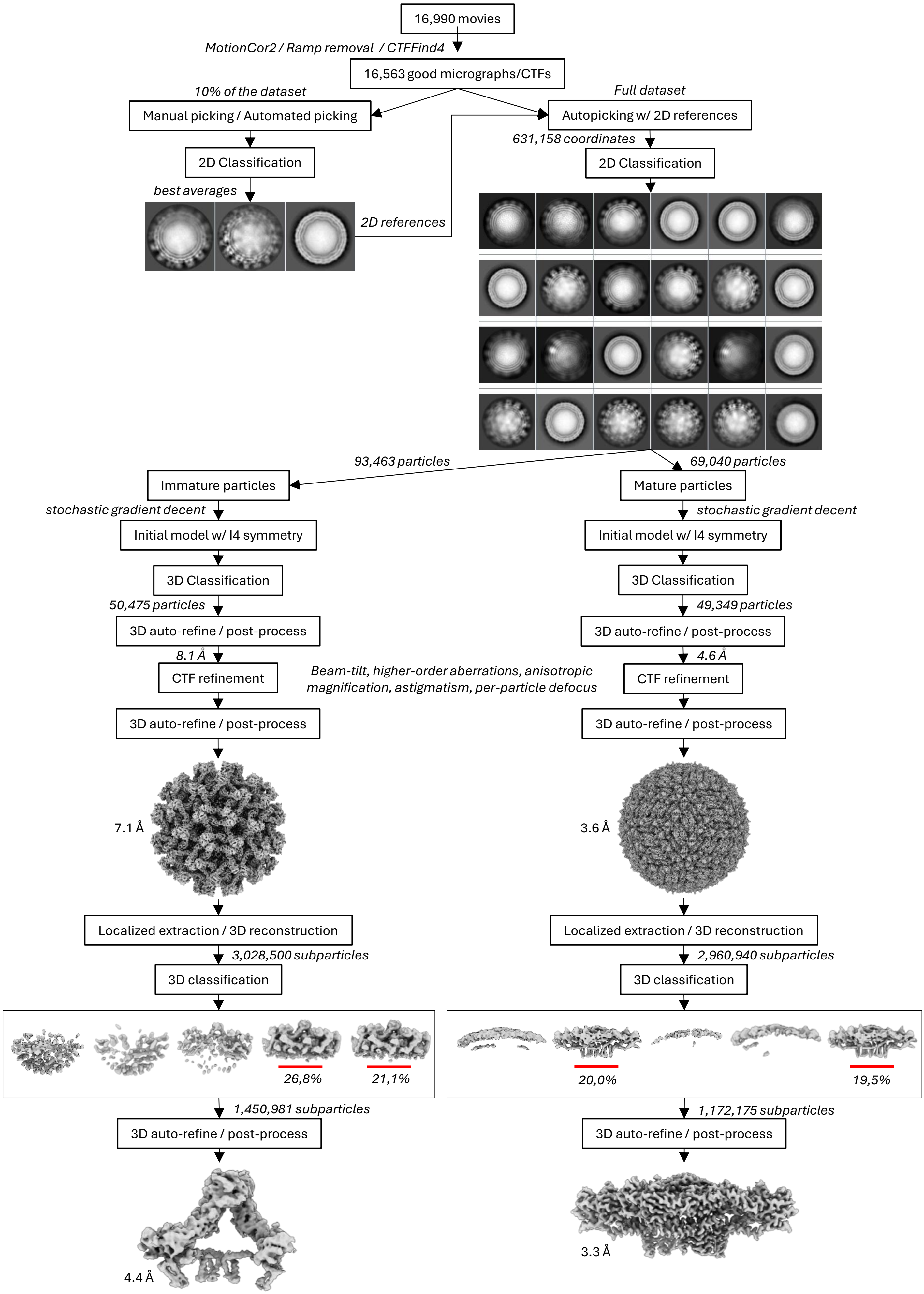


**Supplementary Figure 3 – Single-particle analysis and localized reconstruction workflow used for immature and mature SLEV particles.**

**Supplementary Figure 3 – Single-particle analysis and localized reconstruction workflow used for immature and mature SLEV particles.** Representative scheme of cryo-EM image processing performed within *SCIPION 3*. A total of 16,990 movies were subjected to motion correction, background subtraction, and CTF estimation, yielding 16,563 micrographs suitable for further analysis. Initial manual and automated particle picking performed on 10% of the dataset generated representative 2D class averages that were subsequently used as templates for reference-based autopicking across the full dataset. After 2D classification, mature and immature particles were separated based on morphology and processed independently. Successive rounds of initial model generation, 3D classification, auto-refinement, post-processing, and CTF refinement yielded final full-particle reconstructions at 7.1 Å (immature) and 3.6 Å (mature) of global resolutions. For localized reconstruction, asymmetric-unit coordinates were defined in *UCSF Chimera 1.19* and used for sub-particle extraction with *LocalRec*. Focused 3D classification identified subsets displaying improved structural features (representative selected classes highlighted by red underlines), which were subjected to additional refinement and post-processing. Final localized reconstructions reached global resolutions of 4.4 Å for the immature ASU and 3.3 Å for the mature ASU. Representative 2D class averages, intermediate maps, particle numbers, and resolutions obtained at each stage are shown throughout the workflow.

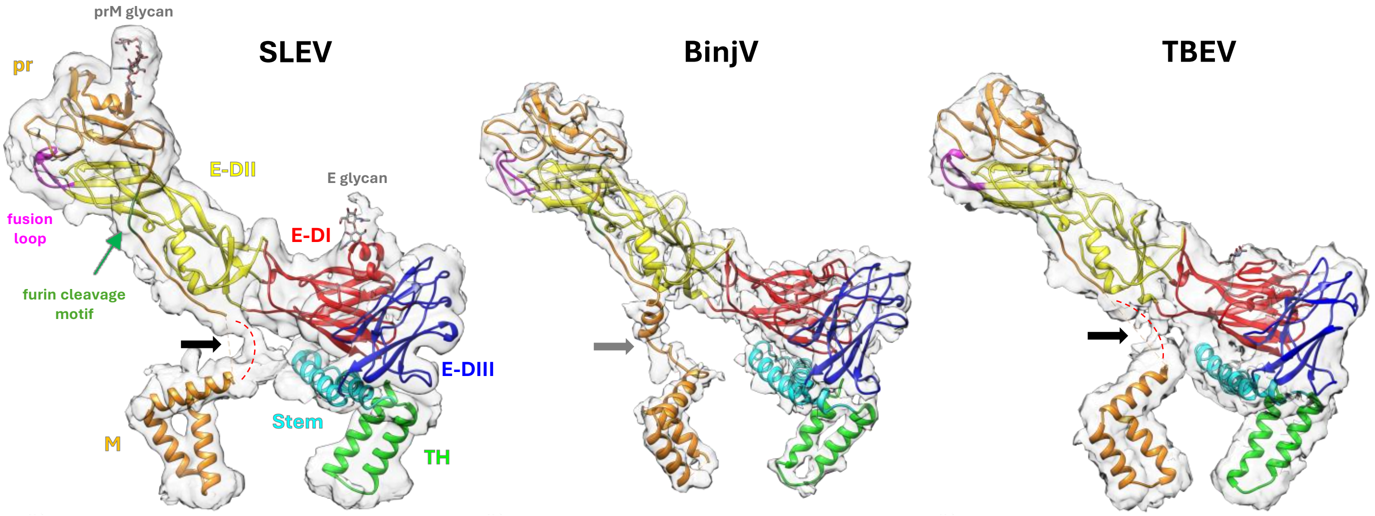


**Supplementary Figure 4 – Comparison of the prM linker region in immature SLEV, BinJV, and TBEV.** Individual prM-E heterodimers from immature SLEV (left), BinJV (middle), and TBEV (right) shown in similar orientations. In SLEV and TBEV, the linker connecting the pr and M domains is partially unresolved, resulting in a gap near the base of the spike. At lower density thresholds, a density bridge can be observed across this region in both structures (black arrows). In BinJV, the corresponding linker is fully resolved and occupies a similar central position beneath E-DII (grey arrow). The trajectories inferred for the unresolved linker segments in SLEV and TBEV (red dashed curves) are consistent with the linker path observed in BinJV, although minor differences in the positions of the M-domain helices may influence the exact path adopted by the linker. Cryo-EM density is shown as a transparent gray surface. E domains and prM are colored as follows: E-DI (red), E-DII (yellow), E-DIII (blue), Stem (cyan), transmembrane helices (TH, lime green), fusion loop (magenta), and prM (orange); the furin cleavage motif is colored in green. The same color scheme is used for the respective regions in the BinJV and TBEV models. BinJV – PDB: 7L30, EMDB: 23147. TBEV – PDB: 8PUV, EMDB: 17946.

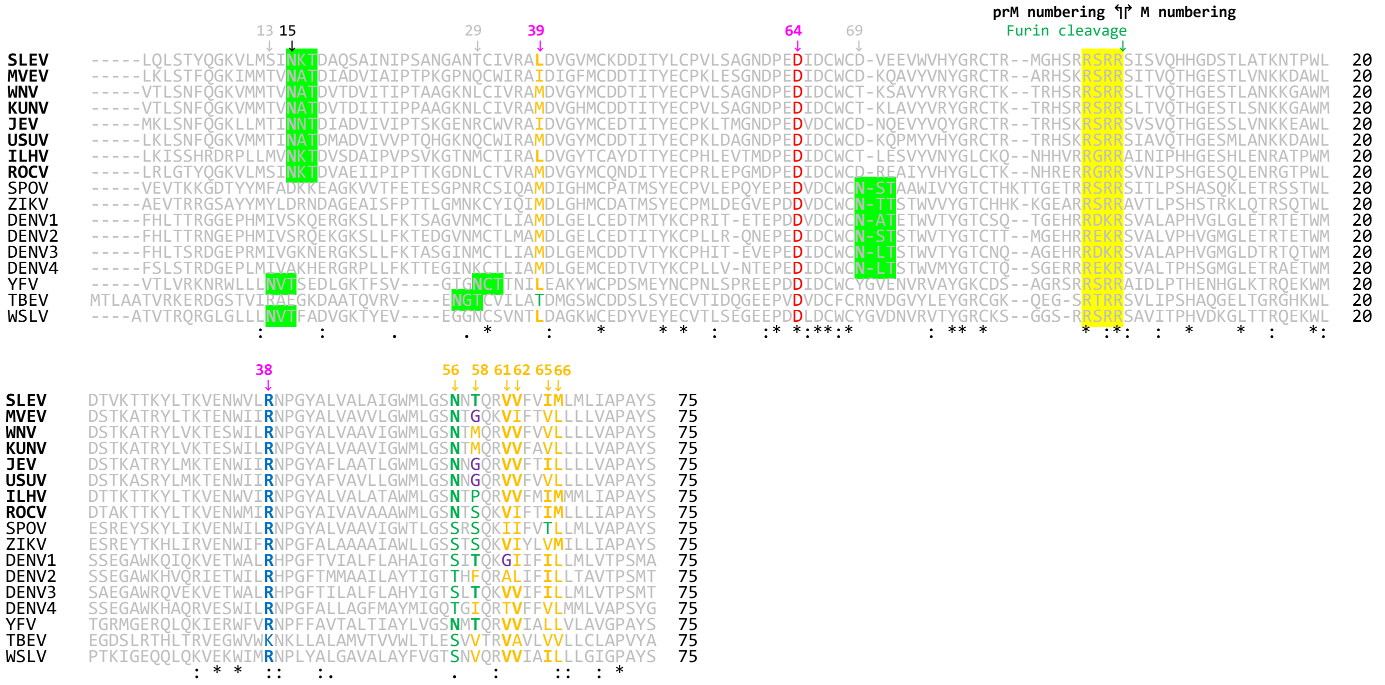


**Supplementary Figure 5 – Conservation of residues in orthoflavivirus prM/M proteins**. Sequence alignment of prM/M proteins from reference sequences of human-infecting orthoflaviviruses generated with Clustal Omega, highlighting the conservation of residues discussed throughout this study and their structural context in SLEV. Residues are colored according to their physicochemical properties: positively charged residues Arg/Lys (dark blue); negatively charged residues Asp/Glu (red); polar residues (green); apolar residues (gold); and glycine residues (purple). Predicted N-linked glycosylation motifs (N-X-S/T) are highlighted by green boxes, and predicted furin cleavage motifs (R-X-R/K-R↓) by yellow boxes. Residue numbering before the furin cleavage site corresponds to prM, whereas numbering after cleavage corresponds to M. The color of residue numbers indicates participation in structural interaction networks identified in SLEV. Gold numbers denote residues associated with conserved lipid-binding pockets. Pink numbers indicate residues involved in E-(pr)M interactions associated with H219 and H246*. The N-linked glycosylation site in SLEV prM occurs at N15 (black number), whereas alternative predicted N-linked glycosylation sites in other orthoflaviviruses are indicated by grey numbers corresponding to their respective residue numbering. Residues shown in bold correspond to interactions mediated primarily by side-chain atoms in SLEV. Virus names shown in bold correspond to encephalitogenic orthoflaviviruses from the JEV serogroup. NCBI RefSeq accession numbers used in the alignment: SLEV (St. Louis Encephalitis Virus, NC_007580.2), MVEV (Murray Valley Encephalitis Virus, NC_000943.1), WNV (West Nile Virus, NC_009942.1), KUNV (Kunjin Virus, D00246.1), JEV (Japanese Encephalitis Virus, NC_001437.1), USUV (Usutu Virus, NC_006551.1), ILHV (Ilheus Virus, NC_009028.2), ROCV (Rocio Virus, NC_040776.1), SPOV (Spondweni Virus, MG182017.2), ZIKV (Zika Virus, NC_035889.1), DENV1 (Dengue Virus type 1, NC_001477.1), DENV2 (Dengue Virus type 2, NC_001474.2), DENV3 (Dengue Virus type 3, NC_001475.2), DENV4 (Dengue Virus type 4, NC_002640.1), YFV (Yellow Fever Virus, NC_002031.1), TBEV (Tick-Borne Encephalitis Virus, NC_001672.1), and WSLV (Wesselsbron Virus, NC_012735.1).

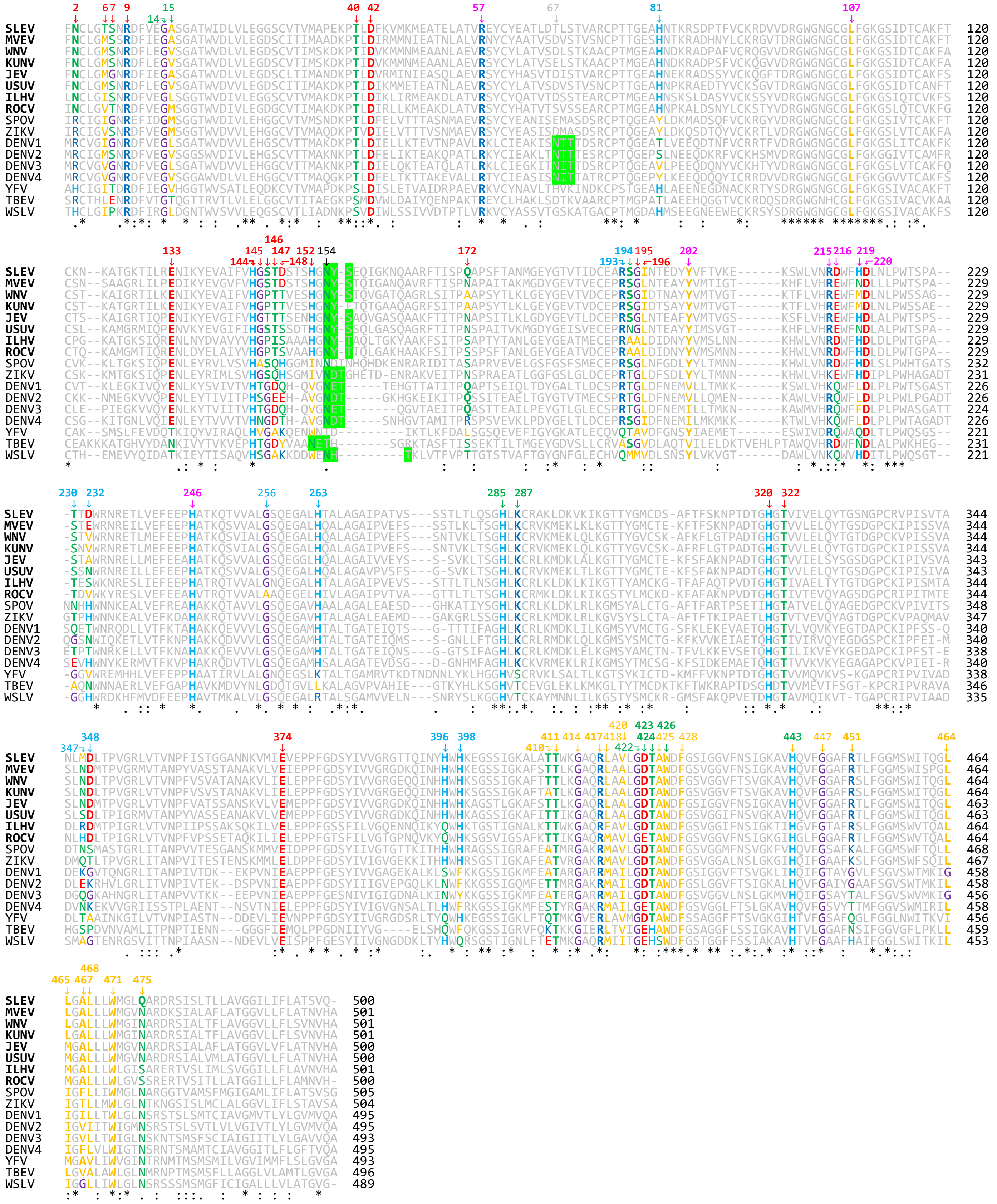

**Supplementary Figure 6 - Conservation of residues in orthoflavivirus E proteins.**

**Supplementary Figure 6 - Conservation of residues in orthoflavivirus E proteins.** Sequence alignment of E proteins from reference sequences of human-infecting orthoflaviviruses generated with Clustal Omega, highlighting the conservation of residues discussed throughout this study and their structural context in SLEV. Residues are colored according to their physicochemical properties: His (light blue); positively charged residues Arg/Lys (dark blue); negatively charged residues Asp/Glu (red); polar residues (green); apolar residues (gold); and glycine residues (purple). Predicted N-linked glycosylation motifs (N-X-S/T) are highlighted by green boxes. The N-linked glycosylation site in SLEV E occurs at N154 (black number), whereas alternative predicted N-linked glycosylation sites in DENV1-4 are indicated by grey numbers. The color of residue numbers indicate participation in the interaction networks identified in SLEV. Gold numbers denote residues associated with the conserved lipid-binding pockets. Red numbers indicate residues involved in the E-DI/E-DIII hinge network associated with H144*, H152, and H320*. Light blue numbers identify residues participating in intra- and interdimer interactions involving H81, H263, H396, and H398. Green numbers denote residues forming the E-DI/Stem interaction network associated with H285* and H443*. Pink numbers indicate residues involved in E-(pr)M interactions associated with H219 and H246*. Residues shown in bold correspond to interactions mediated primarily by side-chain atoms in SLEV. Virus names shown in bold correspond to encephalitogenic orthoflaviviruses from the JEV serogroup. NCBI RefSeq accession numbers used in the alignment: SLEV (St. Louis Encephalitis Virus, NC_007580.2), MVEV (Murray Valley Encephalitis Virus, NC_000943.1), WNV (West Nile Virus, NC_009942.1), KUNV (Kunjin Virus, D00246.1), JEV (Japanese Encephalitis Virus, NC_001437.1), USUV (Usutu Virus, NC_006551.1), ILHV (Ilheus Virus, NC_009028.2), ROCV (Rocio Virus, NC_040776.1), SPOV (Spondweni Virus, MG182017.2), ZIKV (Zika Virus, NC_035889.1), DENV1 (Dengue Virus type 1, NC_001477.1), DENV2 (Dengue Virus type 2, NC_001474.2), DENV3 (Dengue Virus type 3, NC_001475.2), DENV4 (Dengue Virus type 4, NC_002640.1), YFV (Yellow Fever Virus, NC_002031.1), TBEV (Tick-Borne Encephalitis Virus, NC_001672.1), and WSLV (Wesselsbron Virus, NC_012735.1).

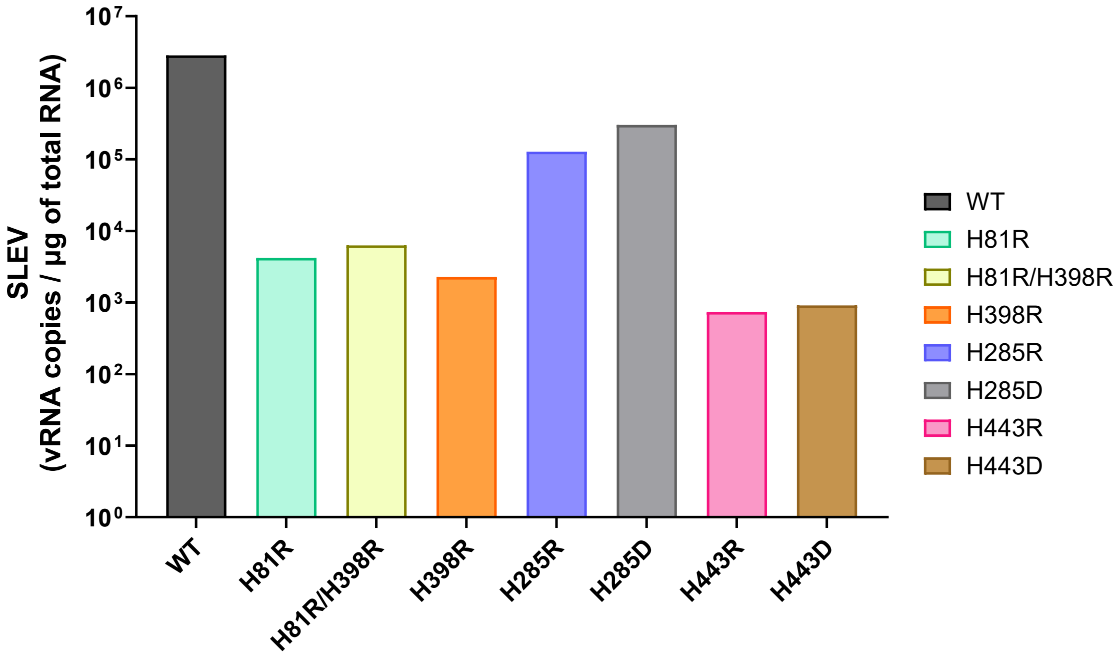


**Supplementary Figure 7 - Recombinant SLEV constructions produce detectable viral RNA in supernatant of HEK293T.** Quantification of SLEV RNA in culture supernatants 8 days post transfection of HEK193T cells with infectious cDNA clones containing selected mutations. Viral RNA levels were quantified by RT-qPCR using SLEV specific primers. Data represent the mean of two technical replicates. Supernatant RNA was treated with DNase to eliminate residual transfected DNA.
